# Beyond Correlation: An Ultra-Large Physics-Driven Vascularized Tumor Model to Bridge Feature Formation with Underlying Biology

**DOI:** 10.1101/2025.11.26.690767

**Authors:** Jiayi Du, Yu Zhou, Lihua Jin, Ke Sheng

**Affiliations:** Department of Radiation Oncology, University of California, San Francisco, San Francisco, CA, USA; Department of Mechanical & Aerospace Engineering, University of California, Los Angeles, Los Angeles, CA, USA

## Abstract

Radiomics provides an appealing, non-invasive approach to probing tumor biology for potential diagnostic and prognostic applications. However, its clinical adoption is limited by challenges in interpretability, which in turn compromise its robustness. To uncover the underlying causation, we developed an ultra-large-scale (ULS) computational model that simulates heterogeneous, vascularized tumor growth under physical constraints to a scale that can be visualized in medical images. Our study revealed the pivotal role of tumor proliferation rate in driving necrosis and tissue heterogeneity and the dominant impact of oxygen consumption rate on vascularization level. Analysis of the resultant tumor Radiomics shows a causal relationship between tumor biophysical parameters and imaging features. Specifically, differences in proliferation and oxygen consumption rates result in distinct changes in radiomic image features, identifying suitable imaging modalities and quantitative imaging metrics for studying these biophysical parameters. This work thus reverse-engineers the building blocks of Radiomics as a means to understand their respective biological underpinnings.

---

Medical images (CT, MRI, PET, etc.) contain far more information than what is visually interpretable to humans, offering untapped potential for disease diagnosis and prognosis^1^. Radiomics leverages high-throughput quantitative features extracted from medical images to build machine learning models that predict clinical outcomes^2^. These features are thought to reflect pathophysiological characteristics, providing non-invasive insights into tumor biology and complementing treatment decisions in personalized medicine. Since the seminal work of Aerts et al.^2^, radiomics has seen exponential growth, with applications in predicting treatment responses^3^, tumor staging^4^, tissue identification^5^, and assessing cancer genetics^6^.

Despite its promise, radiomics faces critical challenges in interpretability and robustness^7^. As a data-driven approach, Radiomics identifies correlations but lacks causal or mechanistic grounding, which are major weaknesses of a clinical tool, particularly when the information is used to inform clinical decisions. Radiomic features are highly sensitive to image acquisition and reconstruction parameters^8^, with many exhibiting poor repeatability even under identical conditions^9^. The high dimensionality of feature sets—where informative features can be obscured by irrelevant variables—exacerbates spurious correlations, overfitting, and compromised generalizability. Various efforts have been made to apply Radiomics in a more interpretable context. Instead of applying the end-to-end model for prediction, many studies are trying to identify the link between Radiomic features and specific biological groundings such as gene expression^10 11^, microvascular density^12^, and tumor metabolism^13^. Other approaches employ interpretable models, such as linear models and decision trees, to determine feature importance. However, these purely *discriminative* strategies are inherently incomplete and incapable of establishing causal mechanisms between biology and imaging features.

In this work, we enhance and expand the concept of Radiomics by developing a *generative* tumor model that sheds light on the physical processes behind tumor growth patterns. While tumors originate from complex genetic and molecular activity at the microscopic level, the patterns we observe in medical images are shaped by larger-scale physical properties and interactions with the surrounding tissue. Computational modeling allows us to isolate and control specific physical factors, making it possible to study how these traits influence tissue development at the mesoscale. By building a scalable simulation framework, we can link defined biophysical characteristics directly to the statistical patterns found in imaging features.

Our model also generates ground truth datasets for various anatomical and functional images at multiple resolutions, with high computational efficiency. This enables us to assess which types of images and features are most informative—without being limited by real-world noise—offering practical insights for improving Radiomics analyses and choosing the most effective imaging techniques. Although real tumor biology involves many variables beyond current modeling capabilities, focusing on the key drivers of heterogeneity provides a powerful new perspective for understanding and advancing Radiomics.

Technically, we developed a large-scale vascularized tumor growth model that captures heterogeneous tumor development while simulating the evolution above the millimeter scale. Our framework explicitly incorporates key biophysical processes driving macroscopic heterogeneity, including stochastic angiogenesis, which generates heterogeneous perfusion and oxygenation patterns, tissue mechanical deformation from growth, and necrotic tissue formation as a function of metabolic stress. To simulate large, coupled vessel-tissue systems efficiently, we implemented a hybrid computational framework as shown in Fig 1(a): vasculature is represented as discrete graphs embedded within a continuum-mechanics-based deformable tissue domain. This separation allows each subsystem to leverage numerical methods best suited to its physics. Meanwhile, various empirical formulae validated against *in vivo* data are introduced in the microcirculation dynamics modeling, enhancing both fidelity and computational efficiency. Key microscopic biophysical parameters were sourced from literature or estimated via physiologically plausible assumptions where data were unavailable; downstream tumor heterogeneity characteristics are solely the result of the system’s evolution entirely under physical constraints. To eliminate spatial bias in vascularized tumor simulations and address the unique challenges associated with dynamic vascular function modeling, we also introduced a novel randomized vasculature initialization strategy along with a corresponding protocol for assigning boundary blood pressures. As a result, our platform enables physiologically unbiased tumor development to sizes previously unattainable, closely mirroring the biophysical vascular properties and tissue growth patterns observed in actual tumors. An example of modeled tumor and vasculature development is illustrated in Fig. 1(b). Starting from a microscopic cluster of tumor cells surrounded by host vasculature, the tumor develops into a multi-millimeter mass with a complex vascular network and a heterogeneous microenvironment, including spatially varying oxygen levels.

**Fig. 1.**
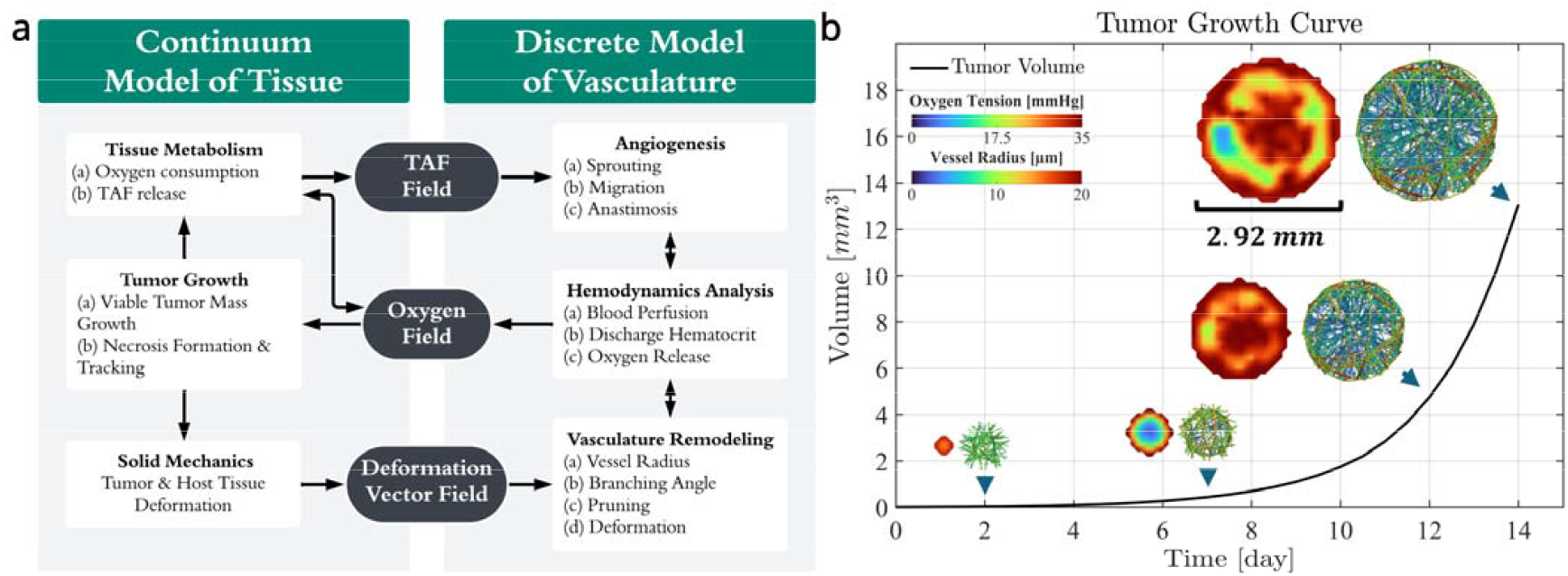
Overview of the tumor model and baseline tumor growth. **(a)** Schematic of the hybrid tumor model, featuring a continuum representation of tissue and a discrete representation of vasculature. White boxes denote simulation modules and their respective functions, while the black boxes highlight the various field data generated and transferred. Arrows indicate the flow of information between modules. **(b)** Growth curve of the baseline tumor over time. The 3D rendered vasculature structure, color-coded by vessel radius, and the central slice tissue oxygen level maps are shown at days 2, 7, 12, and 14 to illustrate structural and environmental changes alongside tumor progression.

By examining the influence of cellular proliferation rate (PR) and oxygen consumption rate (OCR) on tumor patterning and heterogeneity, we have elucidated the mechanistic links between biophysical properties and tumor characteristics. Key findings include the pivotal role of tumor proliferation rate in driving necrosis and tissue heterogeneity and the impact of OCR on tissue vascular density. Using our platform, we generated 20 tumor samples with randomized PR and OCR parameters and attempted to predict their ground truth values using Radiomics features. The resulting high-performance predictive model sees through tumor appearance to identify critical features that uncover underlying biological processes. Given the insight from modeling, the rationale behind feature selection can be understood, and features can be interpreted. We further demonstrated differences in the visibility of these biological processes on imaging features, identifying optimal imaging modalities for specific biophysical insights.

This work establishes a new paradigm by leveraging computational modeling to causally link underlying tumor biology with data-driven imaging features. It offers unique insights into tumor imaging strategies, enabling the selection of modalities tailored to specific biophysical parameters.

## Results

### Model Implementation

In this work, we initialize the system with a spherical tumor embedded at the center of a cubic domain of normal tissue, surrounded by a pre-existing vasculature network. Oxygen supply to the tumor originates either from explicitly modeled blood vessels or through diffusion from the surrounding normal tissue, which is assumed to be supported by an implicitly balanced vasculature. Details of the initialization and boundary conditions are provided in the Supplementary Material S2.2. As the tumor grows, its oxygen demand surpasses local availability, leading to hypoxia and triggering the release of tumor angiogenic factors (TAFs). These factors promote endothelial sprouting and guide the migration of new vascular segments. When two sprouts successfully connect via anastomosis to form a perfused vessel, the new vessel begins supplying oxygen to support further tumor expansion. However, if local oxygen levels fall below a critical threshold before vascularization is established, necrosis occurs in the anoxic regions, where tumor cells lose their metabolic function and can no longer proliferate.

The continuum component of the model, including the reaction-diffusion of diffusive substances such as oxygen and TAF, the growth and elastic deformation of tissue, and the necrosis formation are solved in COMSOL Multiphysics® (v6.1) using its fully coupled solver. Blood perfusion and vasculature development, including angiogenesis and vessel remodeling, are handled in MATLAB® (R2023b) with in-house developed codes, and the communication of field data between the two software is through LiveLink™. The mathematical representation of these models is described in Methods.

### Baseline tumor

The baseline tumor model is calibrated against published data for mouse glioma (GL261), or human glioma data where murine data is unavailable. Starting from 0.0133 mm^3^, the tumor grows exponentially to 13.064 mm^3^ in 14 days (Fig. 1(b)), with the daily growth rate aligning with the fast-growing small *in vivo* gliomas as documented ^14^. Central necrosis does not develop in this scenario due to early vascularization during the initial growth stages that sustain oxygenation. The vascular structure and color-coded maps illustrating the heterogeneous distribution of vessel radius, blood pressure, discharge hematocrit (Hd), blood flow, and wall shear stress (WSS), along with their corresponding histograms, are shown in Fig. 2(a) and 2(c). The internal vascular network morphology is characterized by its complex interconnections and tortuous pathways, including numerous blind ends. The modeled vessel tortuosity is 1.152, closely matching the observed GL261 glioma vasculature tortuosity of 1.15197. Large vascular shunts can be observed connecting surrounding vessels, forming high-flow shortcuts. These shunts are commonly found in tumor vasculature and may impair blood perfusion deeper within the tumor tissue^15^.

**Fig. 2.**
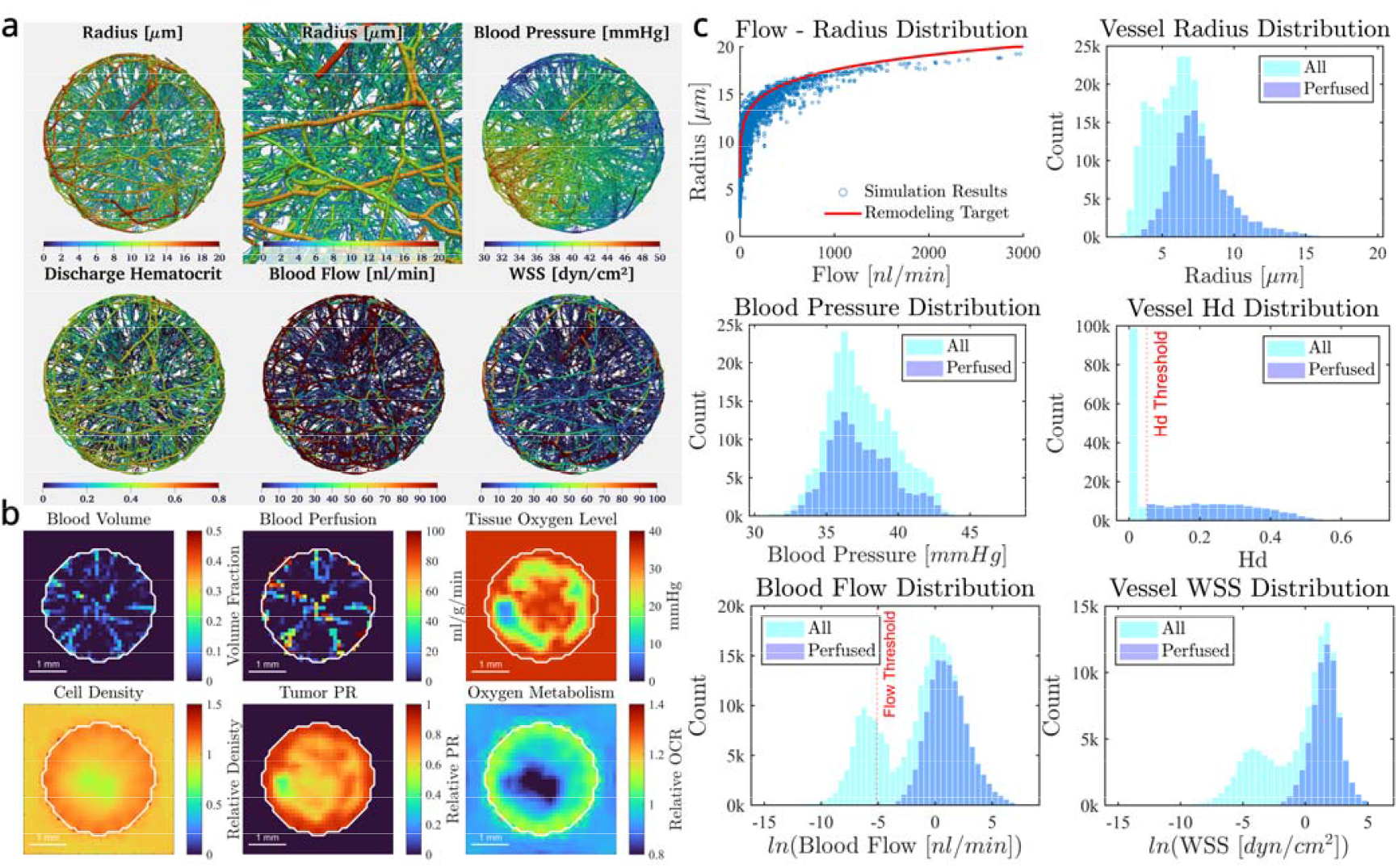
Detailed characterization of the baseline tumor at the end of growth. **(a)** Vasculature morphology with color-coded maps for various vascular properties. From top left to bottom right: (1) vessel radius; (2) zoomed-in vessel radius; (3) blood pressure; (4) discharge hematocrit; (5) blood flow rate; and (6) vessel wall shear stress. **(b)** Central slice maps of key tumor properties, white outline indicates tumor segment boundaries. Displayed example maps include: (1) blood volume; (2) blood perfusion; (3) tissue oxygen level; (4) relative cell density; (5) relative tumor proliferation activity; and (6) relative tissue oxygen metabolism rate. **(c)** Vascular property distributions. (1) Scatter plot of simulated flow–radius relationships overlaid with the target flow–radius curve; (2) histograms of vessel radius; (3) blood pressure; (4) discharge hematocrit; (5) logarithmic blood flow rate; and (6) logarithmic vessel WSS. Perfused vessels are defined as those with both blood flow exceeding 100 µm^3^/s and discharge hematocrit greater than 0.05.

Functional parameters such as blood flow rate and discharge hematocrit—both critical for oxygen delivery—exhibit pronounced heterogeneity, further contributing to the uneven progression of tumor growth. Vessel radius remodeling, guided by WSS, enhances flow transport efficiency by adapting vessel diameter to local flow conditions. As illustrated in Fig. 2(c1), this process leads to a broad distribution of vessel radii ranging from 2 to 20 μm, as shown in Fig. 2(c2). This remodeling ultimately shapes the overall blood flow rate and vessel radius distributions in the assembled GL261 dataset. Quantitative comparisons between simulated tumor characteristics and reference values from the literature are provided in Table 1. All key metrics fall within experimentally reported ranges, validating the model’s biophysical fidelity.

**Table 1.** Quantitative summary of the simulated baseline tumor and the reference value.

|  | Simulation | Unit | Reference |
| --- | --- | --- | --- |
| **Maximum Cell Proliferation Rate** | 1 | $\text{Day}^{-1}$ | |
| **Oxygen Consumption Rate** | 3 | $\text{mmHg/s}$ | 2.2 - 3.7<sup>16</sup> |
| **Growth Time** | 14 | day |  |
| **Tumor Volume** | 13.08 | $\text{mm}^3$ | |
| **Daily Growth Ratio** | 62.88 | % | 13% - 63%<sup>14</sup> |
| **Relative Cell Density** | $1.05 \pm 0.09$ | | |
| **Tissue Oxygen** | $27.68 \pm 6.67$ | $\text{mmHg}$ | 15 - 55<sup>17</sup> |
| **Tissue Perfusion** | $24.22 \pm 108.52$ | $\text{ml/g/min}$ | |
| **Vessel Radius** | $6.46 \pm 2.30$ | $\mu\text{m}$ | 5<sup>18</sup> - 11<sup>19</sup> |
| **Vessel Length Density (perfused / total)** | 79.83 / 136.69 | $\text{mm/mm}^3$ | 76.2<sup>19</sup> - 213<sup>18</sup> |
| **Vessel Surface Density (perfused / total)** | 4.07 / 5.77 | $\text{mm}^2/\text{mm}^3$ | 5.7<sup>19</sup> - 7.5<sup>18</sup> |
| **Vessel Volume Density (perfused / total)** | 1.79 / 2.23 | % | 1.9<sup>18</sup> - 4.8<sup>19</sup> |
| **Blood Flow Rate** | $10.58 \pm 59.33$ | $\text{nl/min}$ | 16.2<sup>19</sup> |
| **Blood Flow Velocity** | $0.44 \pm 1.34$ | $\text{mm/s}$ | 0.44<sup>19</sup> |
| **Wall Shear Stress** | $5.56 \pm 11.68$ | $\text{dyn/cm}^2$ | 17.8<sup>19</sup> |
| **Vessel Tortuosity** | 1.152 |  | 1.151<sup>19</sup> |
| **Branching Length** | $45.49 \pm 55.44$ | $\mu\text{m}$ | 18<sup>18</sup> - 99.7<sup>19</sup> |
| **Bifurcation Density** | 1796.87 | $\text{mm}^{-3}$ | 584<sup>19</sup> - 10000<sup>18</sup> |

Tumor property maps are generated for later analysis, including the tumor and necrosis segmentation, distributions of tissue oxygen partial pressure, cell density, metabolism intensity, hypoxia, proliferation activity, as well as voxel-wise distributions of tissue perfusion and blood volume fraction. The property maps identified for analysis have the potential to be non-invasively imaged *in vivo*. For instance, cell density might be inferred from ADC MRI, which provides information on the extracellular fluid fraction, or from CT scans that reflect density. Tissue blood perfusion and volume could be captured using various imaging modalities with contrast agents, while extracellular oxygen levels might be gauged through Electron Paramagnetic Resonance (EPR)^20^ imaging. Tissue hypoxia could be visualized using 18F-Fluoromisonidazole (FMISO) PET^21^, oxygen metabolism through Oxygen 17 MRI^16^, and proliferation rates through [18F]-FLT-PET^22^. Although these imaging modalities are subject to technological limitations, including low signal-to-noise ratios, coarse resolution, and varying quantitative accuracy, here we consider them ideal imaging methods reflecting the ground truth properties. This approach allows us to focus on identifying the most salient imaging characteristics. Property map examples for the baseline tumor are shown in Fig. 2(b). Detailed definitions of these maps and the potential imaging methodology for each type of information, are all listed in the Supplementary Document S1.2.

### Impact of Growth Parameters Angiogenic Sprouting Rate

Tumor vasculature is a key target in cancer therapy, with interventions taking two divergent approaches. Vascular disrupting focuses on destroying the tumor’s vasculature to indirectly suppress the tumor. Conversely, tumor vasculature normalization aims to improve angiogenesis, thereby enhancing tumor oxygenation, perfusion, and, ultimately, the efficacy of treatments. To explore how different angiogenesis rates affect tumor growth, we conducted simulations on two samples, altering only the sprouting rate from the value in baseline tumor model to 50% and 150% of it, respectively.

Simulation results for the angiogenesis-suppressed tumor reveal a sparsely developed vasculature and a prominent central necrotic core, driven by insufficient blood supply. In contrast, enhancing angiogenesis leads to only modest morphological differences compared to the baseline model. As shown in Fig. 3(a), the tumor with enhanced angiogenesis exhibits the highest growth rate and vascular volume density, while the suppressed model shows the slowest growth. In both baseline and enhanced models, vascular density increases rapidly before reaching a plateau, indicating the formation of a stable vascular network. Meanwhile, the vasculature in the suppressed tumor remains immature and fails to reach equilibrium by the end of the simulation.

**Fig. 3.**
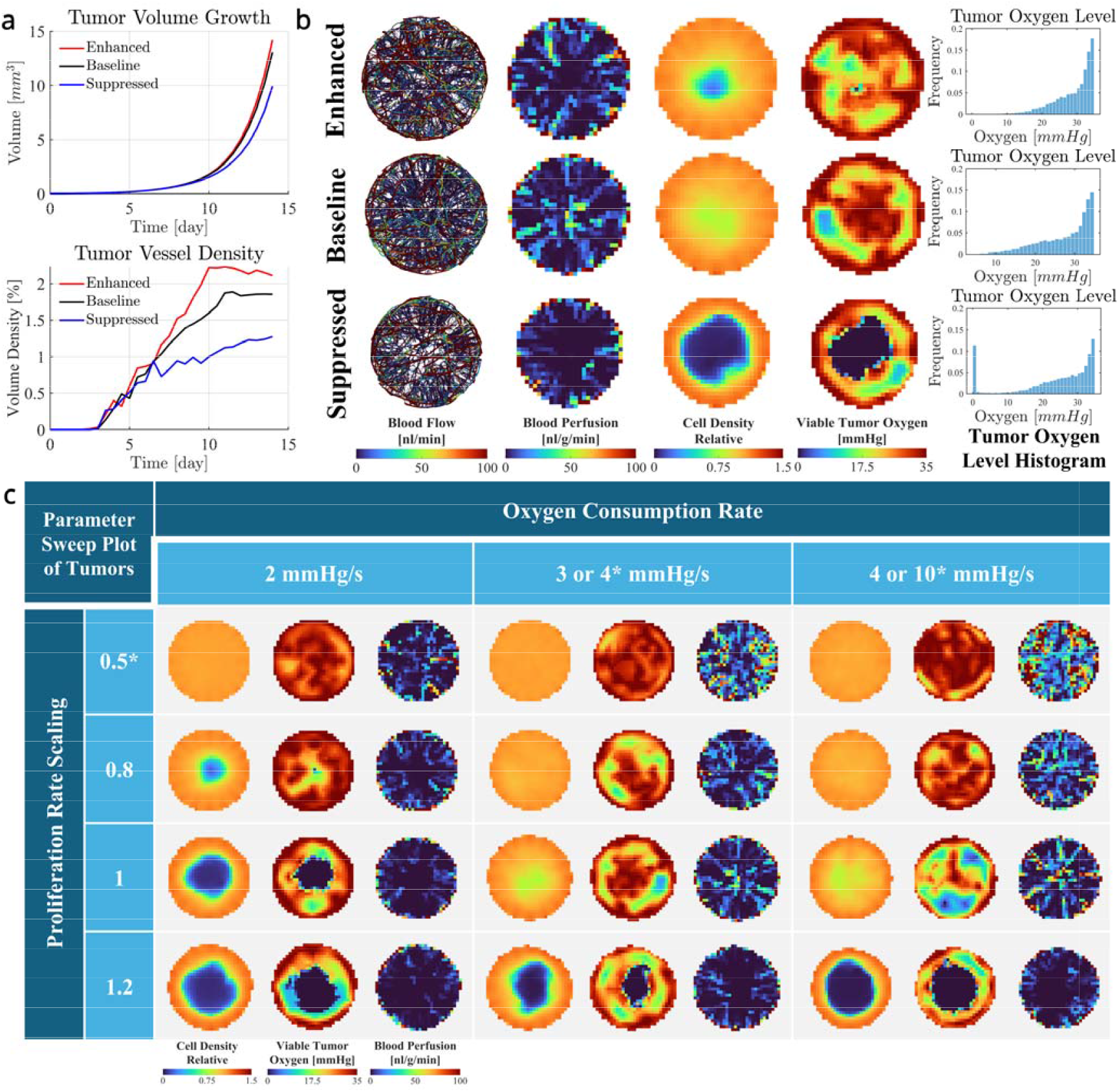
Comparison of tumors with varied growth properties. **(a, b)** Comparison of three types of tumors with different angiogenic sprouting rates: suppressed (-50%), baseline, and enhanced (+50%). **(a)** Volume growth curves and vessel volume density variations over 14 days. **(b)** Tumor characteristics at the end of the 14-day development. From left to right: the first column shows vessels color-coded by blood flow rate, columns 2 to 4 show the tumor’s central slice color-coded by tissue blood perfusion, relative cell density, and oxygen level, respectively, and the final column presents histograms of tissue oxygen level distributions. **(c)** Comparison of tumors with varied oxygen consumption rates and proliferation rates. Analysis time points were selected to ensure similar tumor sizes of approximately 12 mm^3^ across samples. Each cell in the table shows the distributions of blood perfusion, relative cell density, and oxygen level within the tumor central slice. Two groups of tumors were simulated. Slow-growing tumor group*: Proliferation rate scaling of 0.5 with oxygen consumption rates of 2, 4, and 10 mmHg/s. Fast-growing tumor group: Proliferation rate scaling levels of 0.8, 1, and 1.2, combined with oxygen consumption rates of 2, 3, and 4 mmHg/s.

Fig. 3(b) presents spatial maps of vasculature structure, blood perfusion, relative cell density, and oxygen distribution across viable tumor regions. Angiogenesis suppression leads to extensive hypoxic necrosis, as indicated by the spatial oxygen maps. Corresponding oxygenation histograms in Fig. 3(b) show that the baseline and enhanced angiogenesis models exhibit similar oxygen distributions, whereas the suppressed model demonstrates a pronounced shift toward hypoxia. Quantitative summaries, provided in the Supplementary Materials S1.1, reveal strong positive correlations between the angiogenic rate and tissue-level parameters, including oxygenation, perfusion, vessel density, and oxygen uniformity.

The most pronounced contrast is observed between the suppressed and baseline models, while enhancements beyond the baseline result in diminishing returns. This suggests that angiogenic efficacy may saturate in relation to tumor growth demands, potentially limited by autoregulatory mechanisms, such as TAF-mediated feedback on vascular sprouting. As a result, tumors with poor baseline vascularization are likely to respond more robustly to angiogenic interventions, whereas slowly proliferating, well-vascularized tissues—such as benign tumors or normal tissue—may show limited responsiveness. These insights can help guide the strategic use of vascular-targeted therapies, including vascular-disrupting agents and normalization approaches.

### Metabolism and Proliferation

To assess the effects of tumor growth rate and oxygen demand on development, we performed simulations with a range of parameters for nine fast-growing and three slow-growing tumor samples, and analyzed when the tumor volume reached approximately 12 mm^3^. The baseline proliferation rate corresponds to a 24-hour cell doubling time. For fast-growing tumors, the proliferation rate is scaled up to a maximum of 1.2, equivalent to a 20-hour doubling time, consistent with GL261 cell proliferation rates observed in vitro by Szatmári et al.^23^. At the lower end, a scaling factor of 0.5 yields a daily growth ratio of approximately 30%, which aligns with low-proliferation murine GL261 tumors observed *in vivo*^14^, where the daily growth rate from tumor volumes below 1 mm^3^ to approximately 15 mm^3^ ranges between 24% and 49%. For fast-growing tumors, the oxygen consumption rate (OCR) ranges from 2 to 4 mmHg/s, in line with those measured in high-grade (2.2 mmHg/s) and low-grade (3.7 mmHg/s) gliomas^16^. Slow-growing tumors have OCRs from 2 to 10 mmHg/s, with the higher rates approximating the oxygen demand of normal brain tissue. Fig. 3(c) showcases the parameter selection alongside tumor property maps, including cell density, vasculature volume fraction, and tissue oxygen levels for each sample. Quantitative summaries of these tumors are provided in Tables 2 and 3.

**Table 2.** Growth and tissue-related characteristics of 12 tumor samples.

| Sample Parameters |  | Tumor Growth |  |  | Tissue |  |  |  |
| --- | --- | --- | --- | --- | --- | --- | --- | --- |
| Pro-Scale | OCR<br>[mmHg/s] | Tumor volume<br>[mm<sup>3</sup>] | Daily Growth<br>[%] | Necrosis Volume<br>[mm<sup>3</sup>] | Cell density | Tissue Oxygen<br>[mmHg] | Vas Vol Density<br>[%] | Tissue Perfusion<br>[ml/g/min] |
| 0.5 | 2 | 12.34 | 29.8 | 0 | 1.066±0.011 | 30.94±3.45 | 1.5±3.4 | 14.8±61.4 |
|  | 4 | 11.93 | 29.6 | 0 | 1.066±0.011 | 32.27±2.79 | 3.5±6.0 | 42.5±134.8 |
|  | 10 | 11.93 | 28.9 | 0 | 1.066±0.018 | 30.97±4.28 | 7.2±10.3 | 70.1±168.2 |
| 0.8 | 2 | 12.02 | 48.7 | 0.008 | 1.036±0.147 | 31.0±4.7 | 1.9±8.8 | 23.5±102.7 |
|  | 3 | 11.64 | 50.2 | 0 | 1.065±0.024 | 29.3±5.2 | 2.2±4.4 | 20.5±81.5 |
|  | 4 | 11.05 | 49.7 | 0 | 1.065±0.028 | 29.1±4.8 | 2.8±5.2 | 28.7±97.6 |
| 1 | 2 | 12.38 | 62.2 | 0.889 | 0.950±0.308 | 27.2±9.0 | 1.8±4.1 | 25.0±123.3 |
|  | 3 | 13.08 | 62.9 | 0 | 1.050±0.089 | 27.7±6.7 | 2.2±4.5 | 24.2±108.5 |
|  | 4 | 12.67 | 62.5 | 0.003 | 1.054±0.079 | 25.2±7.7 | 2.5±5.3 | 29.8±111.7 |
| **1.2** | **2** | 13.30 | 77.0 | 2.104 | 0.874±0.386 | 21.7±11.7 | 1.3±3.4 | 19.2±101.5 |
|  | **3** | 13.04 | 76.6 | 1.318 | 0.908±0.356 | 24.9±9.5 | 1.7±4.1 | 22.4±118.2 |
|  | **4** | 11.64 | 75.0 | 2.265 | 0.813±0.443 | 21.8±11.5 | 2.0±4.7 | 22.7±102.8 |

**Table 3.** Vasculature-related characteristics of 12 tumor samples.

| **Pro-Scale** | **OCR**<br>**[mmHg/s]** | **Radius**<br>**[μm]** | **Perfused/Total**<br>**Len Density**<br>**[mm/mm<sup>3</sup>]** | **Blood Flow**<br>**[nl/min]** | **Branching**<br>**Length**<br>**[μm]** | **Bifurcation**<br>**Density**<br>**[mm<sup>-3</sup>]** |
| --- | --- | --- | --- | --- | --- | --- |
| **0.5** | **2** | 6.13±2.76 | 48.1/83.8 | 8.7±39.8 | 55.8±64.7 | 880 |
|  | **4** | 6.77±2.55 | 125.1/195.5 | 12.7±56.8 | 50.0±63.4 | 2407 |
|  | **10** | 7.41±2.33 | 268.6/353.1 | 12.1±57.0 | 37.4±45.2 | 5939 |
| **0.8** | **2** | 6.02±2.71 | 65.1/119.9 | 12.0±60.6 | 61.8±79.22 | 1155 |
|  | **3** | 6.27±2.45 | 80.1/133.5 | 8.8±44.8 | 48.7±60.4 | 1648 |
|  | **4** | 6.58±2.36 | 104.6/162.6 | 10.0±47.3 | 44.9±54.1 | 2193 |
| **1** | **2** | 6.09±2.46 | 59.1/117.0 | 13.2±70.0 | 48.4±57.8 | 1444 |
|  | **3** | 6.46±2.30 | 79.8/136.7 | 10.58±59.3 | 45.5±55.4 | 1797 |
|  | **4** | 6.66±2.44 | 91.0±148.0 | 12.3±57.0 | 38.2±47.7 | 2357 |
| **1.2** | **2** | 5.97±2.40 | 42.8±93.6 | 13.7±74.3 | 48.9±56.2 | 1116 |
|  | **3** | 6.40±2.35 | 61.0±107.4 | 13.6±68.1 | 45.2±52.7 | 1401 |
|  | **4** | 6.72±2.29 | 70.5±115.6 | 13.9±65.8 | 40.6±47.0 | 1699 |

As shown in Fig. 3(c) and Table 2, our simulations indicate that cell proliferation is the dominant factor driving tumor growth, while the tumor’s oxygen consumption rate (OCR) plays a secondary, mildly suppressive role—higher OCRs lead to slightly slower growth. Tumors with rapid proliferation develop pronounced cell density heterogeneity and central necrosis, hallmarks of aggressive behavior. In contrast, slow-growing tumors remain uniform in structure and do not show necrosis, regardless of OCR. Importantly, even in the absence of necrosis, fast-growing tumors exhibit significantly higher variability in cell density, highlighting their intrinsic spatial heterogeneity.

Tumor growth rate has a strong inverse effect on average tissue oxygenation, leading to greater oxygen heterogeneity. In contrast, the influence of oxygen consumption rate (OCR) on these metrics is more modest, likely buffered by self-regulating angiogenic responses that adjust vascular development to meet metabolic demands.

Central necrosis is primarily driven by elevated initial tumor growth rates. Among tumors with moderate proliferation rates (scaled at 0.8 and 1.0), those with lower oxygen consumption develop necrosis, while those with higher OCR do not—likely due to slightly higher effective proliferation in low-OCR tumors. This aligns with *in vivo* patterns seen in glioblastomas, where rapidly growing, lower-OCR high-grade GBMs often exhibit necrosis, unlike their slower-growing, higher-OCR low-grade counterparts. At higher proliferation rates (1.2 scale), necrosis occurs regardless of OCR. The severity of necrosis correlates with both growth rate and oxygen demand, with elevated OCR steepening the oxygen gradient and narrowing the viable tissue rim.

As shown in Table 3, vascularization is primarily driven by the tumor’s OCR, as higher oxygen demand stimulates greater vessel recruitment via increased release of tumor angiogenic factors (TAFs) to support growth. Quantitative measures of vascular density—such as vessel count and bifurcation density—show an approximately linear correlation with OCR. In contrast, average functional and other morphological properties of the vasculature, including mean blood flow rate, vessel radius, and branching length, increase only modestly with rising OCR. This suggests that while OCR has a significant influence on vascular network density, its impact on individual vessel characteristics is more limited and complex.

### Radiomics Analysis of Random Tumor Samples

While the tables of semantic tumor characteristics illustrate differences among tumor types based on selected differentiation parameters, such handcrafted metrics represent only a narrow view of the complex spatial and structural patterns that emerge during tumor growth. This prompts a compelling question: Can data-driven, agnostic features—designed to quantify heterogeneity—more effectively reflect variations in underlying tumor biophysics? Conversely, can our mechanistic, physics-informed models enhance radiomics by providing robust, causally grounded insights into critical tumor attributes?

To explore these possibilities, we generated 20 random examples with proliferation rate scaling from 0.8 to 1.2 and oxygen consumption rates from 2 to 4 mmHg/s. Analyses are performed when the tumor volume reaches approximately 12 mm^3^. For each sample, we generated a set of 3941 features, comprising 21 semantic features characterizing vasculature morphology and function, and 3920 predefined agnostic features extracted from tumor maps using PyRadiomics^24^, mainly characterizing tumor heterogeneities. These agnostic features include 14 tumor shape features and 558 quantitative features for each of the seven tumor maps. These maps include relative cell density, oxygen partial pressure, cell oxygen metabolism rate, hypoxia tracer binding rate, vascular volume fraction, blood perfusion, and proliferation activity maps. The extensive feature set for each map encompasses various features in quantitative analyses—first-order statistics, gray level co-occurrence matrix (GLCM), gray level dependence matrix (GLDM), gray level run length matrix (GLRLM), gray level size zone matrix (GLSZM), and neighboring gray-tone difference matrix (NGTDM)—enhanced by different filters.

### Correlation with Radiomic Features

To evaluate associations between radiomic features and tumor proliferation rate (PR) or OCR, we computed the correlation coefficients. Features with absolute correlations >0.5 were classified as strongly linked to these properties. Given the small sample size and high feature dimensionality—a scenario prone to spurious correlations—we performed 100 iterations of correlation analysis between features and randomly generated synthetic properties, establishing a null distribution for false discovery estimation.

The correlation distributions for actual tumor properties (PR, OCR) versus random properties diverge markedly (Fig. 4a). For PR, numerous features exhibit correlations >0.9, indicating strong ties to proliferative activity. In contrast, most features show weak correlations with OCR. For randomly generated values, in contrast to the PR/OCR cases, feature correlations cluster near zero, with virtually no absolute values exceeding 0.7. This confirms that the strong correlations observed with PR and OCR are unlikely to have occurred by chance.

**Fig. 4.**
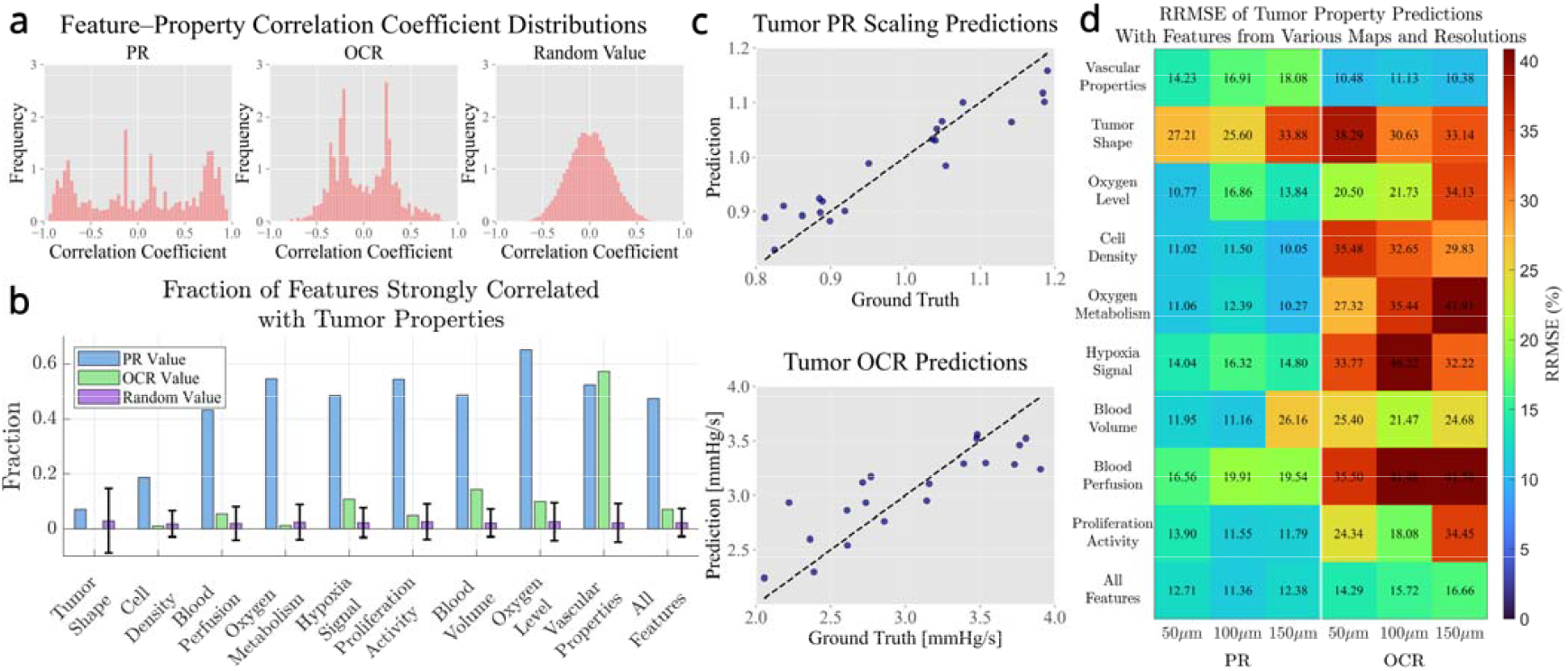
Tumor feature analysis from random tumor samples. **(a)** Distributions of correlation coefficients between extracted tumor features and three variables: tumor proliferation rate scaling factor (PR) (left), oxygen consumption rate (OCR) (middle), and a random meaningless number (RAND) assigned to tumor samples (right). **(b)** Fraction of features extracted from each property map that are strongly correlated with PR, OCR, or RAND. **(c)** Prediction performance for PR and OCR using all features with a LASSO regression algorithm. **(d)** Heatmap of the relative root mean squared error (RRMSE) for LASSO predictions based on features extracted exclusively from specific property map types and resolutions.

Notably, the number of features that strongly correlated with PR far exceeded those associated with OCR. Features extracted from tissue the oxygenation/hypoxia maps and relative proliferation intensity maps show the strongest correlations with PR, whereas features from blood volume and perfusion maps are more closely linked to OCR (Fig. 4b). Importantly, the quantity of features correlated with PR or OCR is substantially greater than those correlated with randomly assigned properties, indicating that these associations likely reflect genuine biophysical relationships rather than random statistical variation.

### Lasso Prediction with Specific Property Map

To predict the two key biophysical parameters—proliferation rate (PR) and oxygen consumption rate (OCR)—we applied Lasso regression, leveraging the magnitude of the resulting coefficients to assess feature importance. All features were standardized prior to training, and the leave-one-out cross-validation was used to enhance model robustness and reduce overfitting. As shown in Fig. 4c, the models achieved relative root mean squared errors (RRMSE) of 11.36% for PR and 15.27% for OCR, significantly outperforming sanity check models. For comparison, a mean-value dummy regressor yielded an RRMSE of 28.9%, and a uniform random predictor returned 40.8%. These results indicate that the radiomic features effectively capture biologically relevant variation in tumor dynamics.

Moreover, the selected features align well with known spatial patterning linked to PR and OCR, offering interpretable connections between imaging features and underlying tumor physiology.

For PR prediction, the most predictive feature is the 90th percentile of the original cell density map, reflecting the model’s finding that higher proliferation rates generate a denser viable rim at the tumor periphery. The second-ranked feature is the Informational Measure of Correlation (IMC2) from the Gray Level Co-occurrence Matrix (GLCM), computed on a high-pass wavelet-filtered cell density map. Lower IMC2 values suggest greater texture complexity or irregularity, which may indicate larger necrotic regions commonly seen in highly proliferative tumors. The third key feature is the complexity metric from the Neighboring Gray Tone Difference Matrix (NGTDM), derived from a high-pass filtered hypoxia map. Lower values of this feature correspond to smoother and more homogeneous hypoxia textures, often associated with widespread necrosis in rapidly growing tumors.

The most significant feature for OCR prediction is the perfused vessel length fraction, which correlates with increased vessel density and connectivity driven by higher metabolic demands, as shown in Table 3. The second most important feature is the small-area low gray-level emphasis (SALGLE) from the Gray Level Size Zone Matrix (GLSZM), calculated on a high-pass wavelet-filtered blood volume map. This feature highlights small, low-intensity regions and may be associated with the reduced extent of avascular regions in high-OCR tumors, as illustrated in Fig. 3c. The third feature is the 90th percentile of the Laplacian of Gaussian–filtered tissue oxygen map, which captures sharp spatial gradients in oxygen levels. This feature negatively correlates with OCR, as tumors with lower OCR are more likely to develop necrotic cores that exhibit steep oxygen gradients.

To assess the predictive value of different tumor property maps and the effect of imaging resolution, Lasso regression was conducted using features exclusively extracted from individual property maps at various resolutions. The resulting relative root mean squared errors (RRMSE), shown in Fig. 4d, indicate that PR was more consistently predicted across property maps compared to OCR, underscoring its stronger influence on tumor spatial patterning. This suggests that Radiomics may be more effective in identifying PR-related mutations than those associated with OCR. Among the property maps, cell density and oxygen metabolism maps were the most effective for predicting PR, while vasculature properties and blood volume maps provided better predictions for OCR, which generally benefits from higher resolution imaging. Notably, using all features together resulted in poorer model performance than selecting the most relevant features from specific property maps.

This highlights the importance of choosing appropriate imaging modalities and feature sources to optimize Radiomics-based prediction.

## Methods

To model the heterogeneous tumor growth with adequate emphasis on the heterogeneity in tumor oxygenation and cellular density while controlling computational costs, we adopted a hybrid approach, which combines a continuum model for averaged cell behavior in tissue with a discrete model handling the perfusion and development of vascular systems. The continuous tissue and discrete vasculature are coupled so that the tissue mechanically deforms the vasculature as it grows and controls the angiogenesis process through the release of TAF, and the vasculature determines the nutrient supply across the tumorous region, which promotes the growth of the tissue. In the study, the healthy mouse brain is the reference site for the parameters of the healthy host; as for the tumor, mouse glioma (GL261) is the reference type. If available, we will prioritize adopting literature-reported parameters for these specific sites. The following section details the construction of the tissue continuum, biochemical, and vasculature models.

### Continuum mechanics model of tissue

The constitutive behavior of both tumor and normal tissue is assumed to be hyperplastic. The strain energy density per grown volume 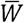 is described by the Blatz-Ko free energy function. The model characterizes the porous, foam-like rubber materials and is widely used for describing the compressible and nonlinear behavior of tissue and tumor^25^:

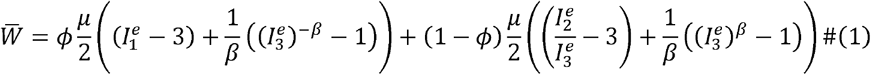

Where *μ* is the shear modulus, 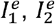, and 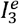 are the invariants of the elastic strain tensor *C*^*e*^ = ***F***^*eT*^*F*^*e*^, given by 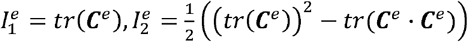, and 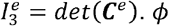. and *β* are non-dimensional parameters, lower *ϕ* lead to more foam-like behavior and *β* is related to the Poisson’s ratio through ν = *β* /(1+ 2 β). While tumors are generally stiffer than normal tissue^26^, there are contradictory reports regarding glioblastoma^27^. In this work, we primarily reference Richard Moran et al.^28^ and James MacLaurin^25^ for the mechanical properties of the normal brain and glioblastoma, respectively. The relative cell density *n*_c_ is estimated as the inverse Jacobian of the elastic deformation gradient *J*_*e*_ =*det*(*F*^e^).

The strain energy density per initial volume *W*is related to 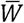 through 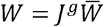. The first Piola-Kirchhoff stress is defined as 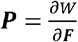, and the Cauchy stress is related to the first Piola-Kirchhoff stress via 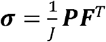.

The tissue is assumed to be under mechanical equilibrium and quasi-static deformation. The balance of linear momentum in the Lagrangian framework is written as

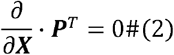

where ***X*** denotes the material point coordinate in the reference configuration.

We assume isotropic growth of the tumor, and the growth tensor is expressed as *F*^*g*^ = *λ*^*g*^ ***I***, where *λ*^*g*^ is the growth stretch ratio that represents the local tumor mass growth. An evolution equation for the growth is given by:

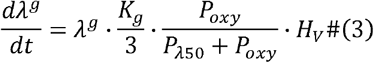

Where *K*_*g*_ is the growth rate parameter, *P*_*oxy*_ is the local oxygen partial pressure, *P*_*λ50*_ is the critical oxygen partial pressure when the growth rate reaches the half maximum. *H*_*v*_ is a cell viability indicator that tracks the irreversible necrosis transition:

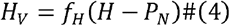

where *H* tracks the lowest oxygen partial pressure experienced by each material point over its growth, and *f*_*H*_ is the Heaviside step function, which returns one if *H* ≥ *P*_N_ and zero elsewise. The tissue will die and stop growing if its goes below the critical oxygen partial pressure for necrosis *P*_*N*_, i.e.*H*_*v*_ becomes zero.

The effective proliferation rate (doubling rate) *r*_*pro*_ of viable cells can be calculated as:

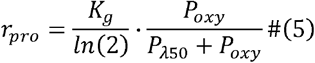

The GL261 population doubling time is measured as 20 hours in vitro^23^. However, for in vivo tumors, the acid environment and insufficient oxygen and glucose supply could dramatically alter the tumor cell proliferation rate^29^. As a result, the typical tumor doubling time is reported to be 2.4 days^30^ with a large variation measured ranging from 1.4 to 6.1 days^14^. Adapting to the reported data, we set the baseline maximum growth rate to have a doubling time of one day.

Parameters for tumor growth and mechanics modeling can be found in the Supplementary Document S2.3.

### Diffusion-reaction of oxygen

Oxygen distribution in homogeneous tissue is governed by the following equation, considering the oxygen diffusion in tissue, the oxygen consumption by tissue, and the oxygen supply through perfused vasculature:

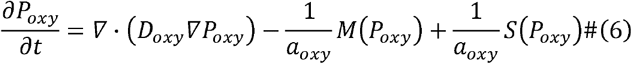

Where *P*_*oxy*_ is the oxygen partial pressure, *D*_*oxy*_ and *a*_*oxy*_ are the oxygen diffusion coefficient and solubility in tissue, respectively. *M(P*_*oxy*_) is the Michaelis-Menten type tissue oxygen consumption, reads:

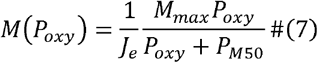

The Jacobian of the elastic deformation gradient *J*_*e*_ quantifies elastic tissue volume change, and its inverse is employed as an approximation for relative cell density. *P*_*M50*_ is the critical oxygen partial pressure when the consumption rate reaches the half maximum, reported to be a low value, typically 0.5 mmHg^31^. *M*_*max*_ is the maximum oxygen consumption rate (OCR), which can vary dramatically across cell types^32^. Tumor tissue metabolism can be significantly lower than that of normal tissue known as Warburg Effect^33^. For human gliomas, the mean OCR has been measured at approximately 2.2 mmHg/s for high-grade tumors and 3.7 mmHg/s for low-grade tumors^16^.

*S* is the volumetric oxygen supply from sufficiently perfused vasculature. The oxygen supply rate of a vessel segment can be written as^34^:

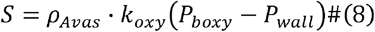

Where *ρ*_*Avas*_ is the surface area density of the vessel segment, *P*_*boxy*_ is the blood oxygen partial pressure and is modeled as constant for perfused vessels, and *P*_*wall*_ is the tissue oxygen partial pressure at the vessel surface. *k*_*oxy*_ is the mass transfer coefficient (MTC) for transvascular oxygen release^34^:

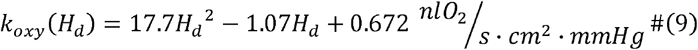

Where *H*_*d*_ stands for discharge hematocrit. In computation, a cross-wall oxygen partial pressure difference cap *P*_*cap*_ is applied to avoid the overestimation of oxygen transportation across the vessel wall due to the limited spatial resolution. In our model, only blood vessels with perfusion higher than 100 μm^3^/s and discharge hematocrit above 0.05 are considered perfused and capable of supporting the tissue oxygen demand; the unperfused vessels are excluded from the oxygen supply.

Despite the high degree of microvascular heterogeneity arising from both intrinsic and extrinsic factors, control mechanisms in healthy tissue effectively buffer fluctuations in oxygen delivery, maintaining relatively stable and adequate oxygen concentrations across various organs^35,36^. Building on these findings, we introduce a dynamic equilibrium host tissue oxygen model. In this approach, the host vasculature is treated as a uniform oxygen source that compensates for tissue oxygen consumption to maintain physiological levels. Oxygen delivery from the explicitly modeled vasculature is restricted to the tumor region. This setup establishes a physiologically relevant boundary condition that allows for the creation of realistic oxygen gradients at the tumor boundary, which are essential for inducing angiogenesis in response to local hypoxia near the tumor-host interface, while simplifying the model by avoiding the need to explicitly simulate healthy tissue vasculature.

### Tumor angiogenic factor

In our model, we introduce a single, unitless, homogenized chemical modulator—referred to as Tumor Angiogenic Factor (TAF)—to represent essential activities needed for the development of vascular networks, particularly those influenced by Vascular Endothelial Growth Factor A (VEGF-A). The concentration of TAF (*C*_*TAF*_) in tissue is governed by:

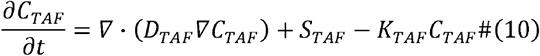

Where *D*_*TAF*_ is the diffusion coefficient of TAF in tissue, *K*_*TAF*_ is the degradation rate. Following JP Alberding et al.^37^ we model the TAF release from hypoxic tissue as:

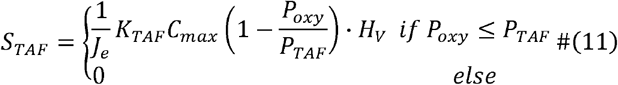

Where *C*_*max*_ is the maximum TAF concentration and *P*_*TAF*_ is the tissue oxygen partial pressure threshold to initiate the TAF release. The viability indicator multiplied in this term reflects the assumption that necrotic tissue is unable to secrete angiogenic factors.

### Vasculature model

In the vasculature model, hemodynamic calculations are performed to estimate blood flow and discharge hematocrit (Hd) distributions. Adequately perfused vessels serve as viable oxygen sources, supplying oxygen to the surrounding tissue. As the tumor grows, the vasculature deforms accordingly, and the inlet and outlet blood pressure conditions are adaptively adjusted to maintain a physiologically plausible pressure gradient and flow within the vascular network. Vascular development is driven by tumor angiogenic factor (TAF), which stimulates the emergence of new sprouts from existing vessels and guides sprout tip migration toward regions of high TAF concentration. These sprouts attempt to anastomose with others, forming new blood flow pathways. The radii of existing vessels also undergo active remodeling to optimize perfusion efficiency—vessels with high flow tend to expand, while those with persistently low flow may shrink or eventually be pruned from the system. The specific components and mechanisms of vasculature modeling are detailed in the sections below.

### Discrete Vasculature Representation

The topology of the vasculature network is described as a multigraph *G* = {*V,E,F*} as a collection of vertices *v*_*i*_ ∈*V*, where vessel segments start and end, edges *e*_*k*_ ∈*E*, which represent the vessel segments between two vertices, and a function *f:E*_⟶{{*vi*_,_*vj*_}: v_*i*,_ *v*_*j*_∈*V and v*_*i*_ ≠*v*_*j*_} that provides a mapping from edges to the vertices they connect. In the discrete vasculature representation, vertices store information such as blood pressure and spatial location, while edges represent vessel radius, blood flow, and discharge hematocrit. Morphologically, this approach captures the full vascular network with minimal computational overhead and enables sub-voxel resolution modeling free from grid artifacts. Functionally, it preserves vascular topology, facilitates efficient flow computation using in vivo–corrected laminar flow assumptions, and directly supports rule-based vascular structure evolution.

### Vasculature Initialization and Boundary Blood Pressure Assignment

The vascularization through angiogenesis requires pre-existing host vasculature. Common methods for establishing host vasculature include the cubic grid vasculature^38^, parallel vessel arrays^39^, and reduced vasculature data containing a few major vessels^40^. However, existing methods fall short in meeting the need for unbiased tumor growth simulations. Artificial vascular arrangements—such as cubic grids or parallel lines—can introduce directional biases, while reduced vasculature density may be insufficient to support nutrient delivery or initiate angiogenesis. Ideally, vasculature extracted directly from normal tissue would be the basis for host vasculature initialization. However, the limited availability of such data, combined with the computational cost of simulating a substantially larger domain, renders this approach impractical. As an alternative, we introduce a spatially stratified tangent vessel initialization method. In this approach, the host vasculature is initialized as tangent lines distributed on a spherical surface at a fixed distance from the tumor boundary. Both the tangent points and vessel orientations are randomly sampled. To prevent unrealistically large vessel-deprived regions, the surface is divided into uniform subregions, and sampling is performed within each to ensure even coverage. This method provides a straightforward yet unbiased vascular environment for tumor development and reduces directional growth bias, thereby enhancing the robustness of statistical analysis.

Accurate assignment of blood pressure at vasculature inlets and outlets is critical for achieving physiologically realistic perfusion and guiding downstream vessel remodeling. In static systems, convex optimization methods incorporating wall shear stress (WSS) under mass conservation constraints have been proposed to estimate boundary pressures^41^. However, such optimization-based approaches are computationally impractical for dynamically evolving vasculature. To address this, we introduce a novel **location-encoded dynamic blood pressure assignment** strategy tailored to our tumor growth model and the specific morphology of the host vasculature. This method incorporates two key components: **length-based pressure gradient term**, which assigns a baseline boundary pressure using estimates of vessel length, reference radius, and target WSS, and ensures a physiologically reasonable pressure gradient along each boundary vessel throughout tumor development, and **orientation-based pressure variation term**, whose component adjusts the boundary pressure based on each vessel’s spatial orientation relative to the tumor center, and which facilitates the formation of realistic pressure gradients within newly formed vessels that connect different host vessels, especially in central tumor regions. Together, these components allow for robust and adaptive pressure assignment throughout tumor growth, maintaining consistent perfusion even as vascular morphology changes significantly. The Supplementary Document S2.2 shows mathematical details of the host vessel initialization and blood pressure assignment.

### Hemodynamics

Assuming Hagen–Poiseuille flow in the lumen of the vessel segment *e*_*k*_ where *f*(*e*_*k*_) = {*v*_*i*_,*v*_*j*_}, the blood flow rate *Q*_*k*_ goes:

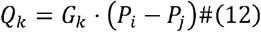

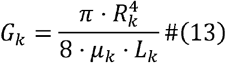

Where *G*_*k*_ is the hydraulic conductance, and μ_*k*_ is the apparent viscosity of the blood flowing inside the vessel. *R*_*k*_ and *L*_*k*_ stands for the radius and the length of the vessel segment. The in vivo apparent blood viscosity is corrected by the Fahraeus-Lindqvist Effect^42^ and endothelial layer (ECL)^43^ effect, both of which are functions of vessel radius and discharge hematocrit.

With the in vivo viscosity and boundary conditions given, a linear system can be constructed by applying mass conservation to all the vertices in the vasculature graph.

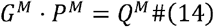

Where *G*^*M*^ is a sparse symmetric matrix for the hydraulic conductance of all vessel segments with

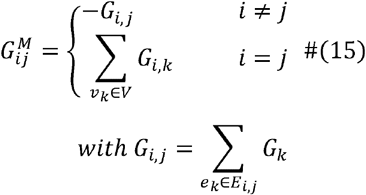

Where *E*_*i,j*_ is the set containing all the edges that connect both vertices *v* _*i*_ and *v* _*j*_. *P*^*M*^ is the vector of blood pressure at vertices, and *Q*^*M*^ is the vector of the blood flow rate of net outflow from vertices. By splitting inner vertex and boundary vertex-related terms into different sides of the equation, the linear system can be transformed to^44^:

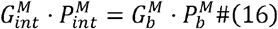

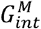 is the submatrix of the hydraulic conductance matrix that contains inner vertex rows, while 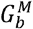 only contains boundary vertex rows. With given boundary blood pressure values 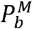, the unknown inner vertex blood pressure 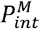 can be effectively solved using the generalized minimum residual method (GMRES)^45^.

Hematocrits have a highly heterogeneous distribution in real vasculature, especially in poorly structured tumor vasculatures^46^, due to the nonproportional distribution of red blood cells (RBCs) in daughter branches at diverging bifurcations, known as the phase separation effect^47^. We adopted an experimentally determined parametric description of the phase separation effect by Pries AR et al.^43^

For computational efficiency, edge contraction is applied to the vasculature graph prior to hemodynamics and RBC calculations, allowing the simulation to focus on key bifurcation and convergence vertices. The blood pressure at the internal vertices along each vessel segment is then derived analytically. This approach accelerates the iterative updates of flow-dependent RBC distribution and RBC-dependent blood flow, which are repeated until convergence. Only vessel segments with sufficient RBC perfusion are considered functional oxygen sources. Details of the empirical formulations used in the hemodynamic calculations are provided in the Supplementary Document S2.1.

### Sprout Activation, Migration, and Anastomosis

The activation of an endothelial cell into a tip cell is modeled as a stochastic process, modified from Alberding et al^37^. The probability of the sprout formation on a vessel segment within a time interval Δ*t* can be written as:

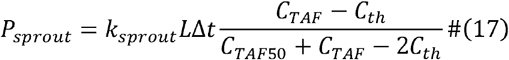

Where *k*_*sprout*_ is the maximal sprout rate per unit length, *L* is the length of the vessel segment, *C*_*th*_ is the TAF concentration threshold for sprout formation, and *C*_*TAF*50_ is the TAF concentration at which the probability of sprouting reaches half-maximum. The new sprout is randomly placed on the activated vessel segment and has a fixed radius *R*_*sprout*_. If the environmental TAF concentration *C*_*TAF*_ is above the tip migration threshold *C*_*mig*_, the stalk cells will be allowed to proliferate, resulting in the sprout elongating at a constant speed *V*_*sprout*_; otherwise, sprout elongation will come to a halt.

The directional sprout elongation is controlled by tip cells, which is influenced by four primary factors: the previous direction, the local TAF gradient, the anastomosis bias, and the random variation. The new migration direction *n*_*new*_ updated from the previous direction *n*_*old*_ goes:

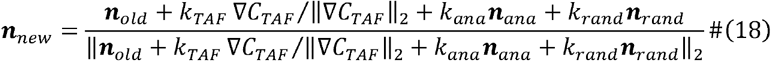

*n*_*ana*_ is the vector indicating anastomosis bias and *n*_*rand*_ is a random 3D unit vector for random variation.*k*_*TAF*_,*kana*, and *k*_*rand*_ serve as the weights for the TAF gradient term, anastomosis bias term, and random variation term, respectively. Adjusted from the work of Secomb et al.^48^, we model the anastomosis bias for an arbitrary tip cell *a* in a form that is both distance- and angle-dependent:

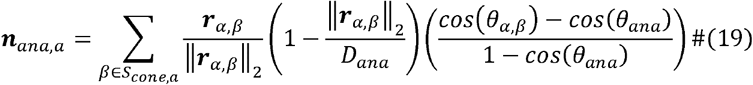

Where *β* refers to other tip cells belonging to the set *S*_*cone,a*_, which includes all the tip cells that can be sensed by tip cell *α*. The vector ***r*** _***α***,***β***_ points from tip cell *α* to tip cell *β*, and *θ*_*α,β*_ represents the angle between the sprout orientation and ***r*** _***α***,***β***_. The tip sensing length limit *D*_*ana*_ reflects the characteristic filopodia length of approximately 75□µm^49^, while the tip sensing angle limit *θ* _*ana*_ captures the forward-extending morphology of filopodia. When two tip cells approach each other within a proximity of the anastomosis threshold *L*_*ana*_, their migration halts, and a vessel segment is formed to connect the two tips, thereby establishing a lumen for potential blood perfusion.

### Radius Remodeling and Pruning

The blood vessel went through radius remodeling to adjust its radius for perfusion efficiency. In this work we propose a novel radius adaptation scheme as follows:

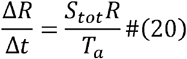

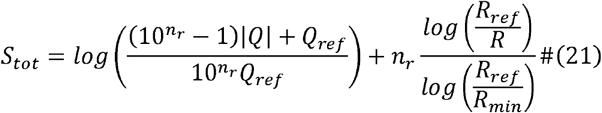

Where *S*_*tot*_ is the total radius remodeling signal, *T*_a_ is the adaptation time controlling the adaptation rate, *n*_*r*_ is a weighting factor for both the target radius and the flow regularization, and it tunes the flow-radius relationship curve. *R*_*ref*_ is the reference radius and *R*_*min*_ is the minimum radius allowed. The reference flow *Q*_*ref*_ is calculated on an individual vessel based on a reference WSS:

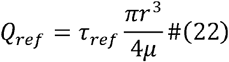

The radius remodeling scheme is designed to halt adaptation once the WSS and vessel radius reach their respective reference values. Vessel radiuses increase when overloaded relative to their current size and decrease otherwise. Zero-flow vessel branches progressively shrink toward a minimum radius *R*_*min*_ and are pruned if the radius falls below a predefined threshold. The update signal scheme is constructed to ensure stability under imposed boundary blood pressure conditions and accommodates various vascular connection architectures, including series, parallel, and hybrid configurations.

Within each vessel update step, the positions of vessel vertices are first updated to reflect tissue deformation. Based on the updated geometry, edge lengths are recalculated, and vessel radii are adjusted according to the previous step’s flow conditions, following equations (20)– (22). Blood pressure at vasculature inlets and outlets is then adapted using the method described in Supplementary Document S2.2. Next, the angiogenesis module simulates sprout formation, tip migration, and anastomosis events, as described by equations (17)– (19). With the updated vascular structure, the hemodynamics module computes blood flow and discharge hematocrit using equations (11)– (16), with additional details provided in Supplementary Document S2.1.

## Discussion

This paper describes a hybrid simulation platform that couples continuum-based tissue dynamics with discrete vasculature modeling, enabling large-scale, hemodynamically detailed simulations of vascularized tumor growth. By incorporating several novel modeling strategies, the platform facilitates unbiased simulations at previously unattainable scales, accurately capturing the biophysical vascular characteristics and growth patterns observed in vivo. We used this framework to investigate tumor progression and spatial organization, establishing a bridge between mechanistic biophysical modeling and data-driven Radiomics. The simulation outcomes provided biologically interpretable Radiomics features and revealed how the detectability of different biophysical parameters varies across functional imaging modalities, emphasizing the critical role of modality selection in Radiomics-driven tumor characterization.

Numerous tumor modeling studies have explored vascularized tumor development. Yet, they often target different objectives and exhibit significant limitations in capturing tumor heterogeneities at the scale we are investigating. For example, Shirinifard^38^ used an agent-based model to show asymmetric tumor cell cluster growth toward vasculature, focusing solely on a submillimeter scale. Similarly, Tobias Duswald et al.^40^ developed a dynamic tree-like agent-based model using BioDynaMo^50^, but their approach omitted crucial processes such as anastomosis in vascularization and assumed that nutrients could be supplied through sprouting blind ends without considering perfusion. Alberding et al.^37,51^ simulated angiogenesis in healthy cerebral and retinal tissues with functional vasculature but restricted their scope to small domains, excluding tumor-induced tissue deformation and growth. Vavourakis et al.^39^ proposed a hybrid model incorporating angiogenesis and perfusion, but it lacked the ability to reproduce tissue heterogeneity, and introduced artifacts, such as overly regular vascular architectures, and growth bias due to its simplified, parallel vessel arrangement.

Our custom ultra-large-scale modeling simulation framework is designed to capture essential real tumor morphological and functional characteristics. The following novel technical approaches enable the framework:

- **Hybrid tissue representation**: The model integrates continuum mechanics for deformable tissue with discrete vasculature modeling, enabling simultaneous simulation of structural deformation and functional vascular dynamics.
- **Empirically grounded microscale modeling**: Empirical rules are extensively employed to represent core vascular functions, ensuring microscale biological fidelity while preserving computational tractability for macroscale simulations.
- **Macroscopic realism with emergent heterogeneity**: The framework captures essential tumor growth dynamics and reproduces realistic vascular morphology and function. Spatial and functional heterogeneities emerge naturally from biophysical interactions, rather than being imposed.
- **Innovative host-tumor interface modeling**: We introduce a novel vasculature initialization protocol and a dynamically equilibrated host tissue environment, which together ensure unbiased tumor expansion and angiogenesis onset without artificial boundary effects.
- **Robust vessel adaptation mechanism**: A newly designed vessel radius adaptation scheme replicates physiological radius distributions and maintains stability under dynamic blood pressure boundary conditions, even in complex vascular networks involving series, parallel, and hybrid connectivity.

Biological systems are intrinsically complex, and modeling them at scale remains an ongoing challenge. While this study introduces a powerful framework for simulating large, heterogeneous tumor growth with unprecedented fidelity, several limitations persist. A major constraint is the limited availability of in vivo data—particularly spatially resolved functional data from individual tumors— which impedes both model calibration and deeper insight into diverse tumor behaviors. Currently, our simulations are confined to mouse-scale tumors, which only partially capture the structural and functional complexity of human-scale malignancies. Computational limitations further restrict tumor size, spatial resolution, and the number of simulations feasible for high-throughput analysis. Our solid mechanics module lacks the ability to model viscoelastic behavior, limiting its accuracy in capturing long-term tissue rearrangement and shape instabilities. Additionally, oxygen source modeling in the vasculature system relies on a threshold-based method that becomes insufficient for tumors exceeding a few centimeters, where oxygen extraction dynamics play a critical role. To address these challenges, future work will focus on enhancing the tissue mechanics model with viscoelastic and plastic deformation capabilities to better represent the evolving tumor architecture. In parallel, we plan to develop high-performance computing platforms to scale simulations to multi-centimeter tumors.

These models will explicitly incorporate intravascular oxygen delivery, enabling more accurate modeling of spatial oxygenation gradients and better alignment with physiological tumor behavior at clinically relevant scales.

The growing availability of RNA sequencing (RNA-seq) data—especially those spatially encoded^52^— opens new avenues for personalized tumor modeling through data-driven parameterization and calibration. By creating a reference database that links gene expression profiles to tumor biophysical properties measured in microscopic-scale experiments, model parameters can be inferred directly from patient-specific RNA-seq data. This approach enables the generation of individualized tumor simulations that reflect each patient’s unique genetic landscape. Moreover, model parameters derived from RNA-seq can be validated and refined using patient-derived organoids, enhancing simulation accuracy. Longitudinal RNA-seq datasets also provide valuable insights into clonal evolution and mutational dynamics, enabling colony-resolved modeling that captures the progression of intratumoral heterogeneity over time.

The model is conducive to supporting the prediction of treatment outcomes and optimization of therapeutic strategies, which was the original promise of Radiomics. Different from genomics, proteomics or transcriptomics that point to potential drug targets, Radiomics demonstrates imaging correlation that cannot be directly targeted. This fundamental limitation and other challenges due to the lack of a mechanistic explanation have prevented Radiomics from being used as a primary instrument for guiding prospective interventions. As cancer therapies advance and diversify, the number of potential treatment combinations has grown dramatically—yet only a small fraction can feasibly be evaluated through clinical trials. Grounded in a robust baseline tumor model and rigorously parameterized biophysical mechanisms, the platform connects microscale responses— shaped by variable genetic and microenvironmental contexts—to macroscale imaging responses. It helps explore key factors such as drug delivery efficiency, vascular remodeling, oxygen enhancement in radiotherapy, and the heterogeneous distribution of both tumor microenvironments and cell colonies. The model can accommodate a wide range of treatment modalities—including radiotherapy, chemotherapy, immunotherapy, receptor-targeted therapy, and vascular-targeted therapy—with flexible support for arbitrary combinations and dosing schedules. By integrating patient-specific biophysical and genetic inputs, the model enables systematic, interpretable exploration of personalized therapeutic strategies for subsequent more effective clinical trials, underscoring its potential as a core tool in advancing precision oncology.

## Code Availability

The source code is freely available and can be found on the GitHub at https://github.com/PhantomOtter/VasTumor.

## Acknowledgements

LJ acknowledges a National Science Foundation MPS-NCI SPARK supplement (No. 2326800) to her CAREER Award (No. 2048219).

## Author information

### Contributions

K.S. and J.D. conceptualized the work. J.D. and Y.Z. designed the methodology. Y.Z. set up the COMSOL simulation. J.D. developed the simulation code, performed analyses, and visualized the data. J.D. wrote the original draft of the manuscript, and all other authors revised it. K.S. and L.J. supervised the work.

