## Supplementary Materials for "Beyond Correlation: An Ultra-Large Physics-Driven Vascularized Tumor Model to Bridge Feature Formation with Underlying Biology"

#### S1 Additional Results

##### S1.1 Impact of Growth Parameters

**Changing Angiogenic Sprouting Rate**

| | | Suppressed | Baseline | Enhanced |
|---|---|---|---|---|
| Tumor Growth | Sprouting Scaling | 0.5 | 1.0 | 1.5 |
| | Growth Time [day] | 14 | 14 | 14 |
| | Tumor Volume [mm³] | 9.91 | 13.08 | 14.22 |
| | Necrosis Volume [mm³] | 1.430 | 0.000 | 0.006 |
| | Daily Growth [%] | 59.7 | 62.9 | 63.9 |
| Tissue Characteristics | Cell density | 0.868 ± 0.396 | 1.050 ± 0.090 | 1.044 ± 0.116 |
| | Tissue Oxygen [mmHg] | 24.0 ± 10.9 | 27.7 ± 6.7 | 29.1 ± 5.8 |
| | Vas Vol Density [%] | 1.6 ± 3.7 | 2.2 ± 4.5 | 2.4 ± 5.3 |
| | Tissue Perfusion [ml/g/min] | 22.8 ± 107.0 | 24.2 ± 108.5 | 34.4 ± 142.5 |
| Vasculature Characteristics | Perfused/Total Length Density [mm/mm³] | 49.3 / 105.6 | 79.8 / 136.7 | 81.4 / 131.2 |
| | Perfused/Total Surface Density [mm²/mm³] | 2.6 / 4.2 | 4.1 / 5.8 | 4.4 / 5.8 |
| | Perfused/Total Volume Density [%] | 1.3 / 1.6 | 1.8 / 2.2 | 2.0 / 2.4 |
| | Blood Flow [nl/min] | 15.4 ± 85.9 | 10.6 ± 59.3 | 15.7 ± 71.8 |
| | Blood Flow Velocity [mm/s] | 0.58 ± 1.89 | 0.44 ± 1.34 | 0.60 ± 1.62 |
| | WSS [dyn/cm²] | 7.0 ± 16.0 | 5.6 ± 11.7 | 7.1 ± 13.5 |

| | | | | |
|---|---|---|---|---|
| | **Branching Length [µm]** | 69.0 ± 76.9 | 45.5 ± 55.4 | 38.2 ± 49.2 |
| | **Bifurcation Density [mm$^{-3}$]** | 860 | 1797 | 2103 |

*Table S1 Summary of tumor samples with suppressed, baseline, and enhanced angiogenesis sprouting.*

#### S1.2 Definition of Property Maps

At the conclusion of each simulation, key tumor development variables are exported from COMSOL to MATLAB and interpolated from tetrahedral finite element mesh onto a Cartesian grid of isotropic voxels (50, 100, or 150 microns) using MATLAB's 'griddata' function. This process prepares the data for in-depth tumor characterization and analysis. Subsequently, comprehensive property maps are generated on this grid and analyzed, covering various aspects such as tumor and necrosis segmentation, distributions of tissue oxygen partial pressure, cell density, metabolism intensity, hypoxia, proliferation activity, as well as voxel-wise distributions of tissue perfusion and blood volume fraction. By analyzing these ground truth maps—unaffected by imaging-related limitations such as signal-to-noise ratio, contrast mechanisms, or resolution constraints—we focus on identify inherently the most informative properties and optimal spatial resolutions for tumor assessment, offering insights to guide future imaging strategies and technology development.

Although the property maps generated for analysis are not meant to mimic medical images but are ground truth directly calculated from the simulation results, these properties have the potential to be non-invasively imaged in vivo. For instance, cell density might be inferred from ADC MRI, which provides information on the extracellular fluid fraction, or from CT scans that reflect atomic composition. Tissue blood perfusion and volume could be captured using various imaging modalities with contrast agents, while extracellular oxygen levels might be gauged through Electron Paramagnetic Resonance (EPR)[1] imaging. Tissue hypoxia could be visualized using 18F-Fluoromisonidazole (FMISO) PET[2], oxygen metabolism through Oxygen-17 MRI[3], and proliferation rates through [18F]-FLT-PET[4]. Table S2 listed possible modalities that could potentially reflect the tumor properties of interest.

| Property Map | Imaging Modality |
|---|---|
| **Cell Density** | CT, ADC MRI |

| | |
|---|---|
| **Oxygen Level** | EPR[1] |
| **Oxygen Metabolism** | Oxygen-17 MRI |
| **Volumetric Proliferation Rate** | [18F]-FLT-PET[4] |
| **Hypoxia** | FMISO[2] |
| **Tissue Perfusion** | Perfusion CT, Perfusion MRI |
| **Blood Volume** | Perfusion CT, Perfusion MRI |

*Table S2 Imaging modalities reflecting tumor properties.*

The cell density map presents a relative estimation of solid tissue mass density, whose voxel intensity is defined as:

$$I_{\rho} = \frac{1}{J_e}$$

The oxygen metabolism map is calculated as the relative volumetric oxygen consumption rate, aligned with the volumetric oxygen consumption rate term in the oxygenation simulation, defined as:

$$I_{OCR} = \frac{1}{J_e}\frac{M_{max}P_{oxy}}{P_{oxy} + P_{M50}}$$

Similarly, the volumetric proliferation activity map measures the proliferation events within a voxel:

$$I_{pro} = \frac{1}{J_e}\frac{P_{oxy}}{P_{\lambda 50} + P_{oxy}} \cdot H_V$$

The hypoxia map in our study is designed to replicate the signal observed in 18F-Fluoromisonidazole (FMISO) PET imaging. While this map reflects tissue oxygen levels, it specifically highlights regions of low oxygenation and deliberately excludes necrotic areas from contributing to the signal. Consequently, it offers distinct insights with a particular emphasis on hypoxic yet viable tissue regions. The binding rate of FMISO as a function of oxygen partial pressure for living cells is given as[5]:

$$k_b(P_{oxy}) = \frac{1}{J_e}\frac{k_{b0}P_{50b}}{P_{oxy} + P_{50b}} \cdot H_V$$

Where $k_{b0}$ is the maximum binding rate at $k_{b0} = 4.5 \times 10^{-4}\ s^{-1}$, and $P_{50b}$ is the oxygen level at the half-maximum binding rate, given as 1.4 mmHg.

### S2 Additional Methods

#### S2.1 Blood Flow Hemodynamics

##### In vivo Blood Viscosity

Modeling blood flow within a given large vasculature poses a significant challenge. The Navier–Stokes equations for the fluid can be difficult and expensive to solve, and the presence of cellular components, which deviate the blood's behavior from the homogeneous fluid, particularly in micro-vasculatures, makes the problem even more complex.

An alternative strategy involves assuming Hagen–Poiseuille flow within each vessel segment and subsequently adjusting the apparent viscosity using empirical formulas derived from experimental data[6]. This approach has gained widespread acceptance and, to the best of our knowledge, remains the only appropriate method for addressing blood flow in large microvascular systems. Therefore, in this study, we embrace this framework for modeling the hemodynamics of vasculature.

Assuming Hagen–Poiseuille flow in the lumen of the vessel segment $e_k$ where $f(e_k) = \{v_i, v_j\}$, the blood flow rate $Q_k$ goes:

$$Q_k = G_k \cdot (P_i - P_j)$$

$$G_k = \frac{\pi \cdot R_k^4}{8 \cdot \mu_k \cdot L_k}$$

Where $G_k$ is the hydraulic conductance, and $\mu_k$ is the apparent viscosity of the blood flowing within. $R_k$ and $L_k$ stands for the radius and the length of the vessel segment. The reference rat blood plasma's apparent viscosity $\mu_0$ measured at 37℃ is 1.05 cP[7]. An additional two-step correction is required to account for various vessel diameters and red blood cell (RBC) concentration conditions and estimate realistic in vivo apparent viscosity in blood vessels.

The first correction is for in vitro viscosity that considers the plasma cell-free layer due to the centering movement of erythrocytes resulting in lower resistance between flow and tube wall, also known as the Fahraeus-Lindqvist Effect[8]. A. R. Pries et al.[9] proposed an empirical correction for such an effect based on in vitro glass tube experiments:

$$\eta_{vitro} = 1 + (\eta_{0.45} - 1) \cdot \frac{(1 - H_d)^C - 1}{(1 - 0.45)^C - 1}$$

$$\eta_{0.45} = 220 \cdot e^{-1.3D} + 3.2 - 2.44 \cdot e^{-0.06D^{0.645}}$$

$$C = (0.8 + e^{-0.075D}) \cdot \left(\frac{1}{1 + 10^{-11} \cdot D^{12}} - 1\right) + \frac{1}{1 + 10^{-11} \cdot D^{12}}$$

Where $D$ is the measured anatomic vessel diameter in microns, $\eta_{vitro}$ is the relative apparent viscosity and $\eta_{0.45}$ is that of rat blood with discharge hematocrit at a level of 0.45. The term discharge hematocrit is defined as the volume fraction of RBCs delivered by the blood flow[10]. The second correction accounts for the endothelial layer (ECL) that observed to substantially increase the in vivo blood viscosity[11]. The effective thickness of the layer $W_{eff}$ can be calculated by adding an asymptotic component $W_{as}$ and a biphasic component $W_{peak}$ with a peak[7]:

$$W_{eff} = W_{as} + W_{peak} \cdot (1 + 1.18 \cdot H_d)$$

$$W_{as} = \begin{cases} 0 & if\ D \leq 2.4 \\ 2.6 \cdot \dfrac{D - 2.4}{D + 100 - 4.8} & if\ D > 2.4 \end{cases}$$

$$W_{peak} = \begin{cases} 0 & if\ D \leq 2.4 \\ 1.1 \cdot \dfrac{D - 2.4}{10.5 - 2.4} & if\ 2.4 < D \leq 10.5 \\ 1.1 \cdot e^{-0.03 \cdot (D - 10.5)} & if\ 10.5 < D \end{cases}$$

Then the effective diameter and in vivo relative apparent blood viscosity $D_{eff}$ reads:

$$D_{eff} = D - 2W_{eff}$$

$$\eta_{vivo} = \eta_{vitro} \cdot \left(\frac{D}{D_{eff}}\right)^4$$

#### Linear System Construction for Blood Flow

With the in vivo viscosity and boundary conditions given, a linear system can be constructed by applying mass conservation to all the vertices in the vasculature graph.

$$G^M \cdot P^M = Q^M$$

Where $G^M$ is a sparse symmetric matrix for hydraulic conductance of all vessel segments with

$$G_{ij}^M = \begin{cases} -G_{i,j} & i \neq j \\ \sum_{v_k \in V} G_{i,k} & i = j \end{cases}$$

$$with\ G_{i,j} = \sum_{e_k \in E_{i,j}} G_k$$

Where $E_{i,j}$ is the set contains all the edges that connect both vertices $v_i$ and $v_j$. $P^M$ is the vector of blood pressure at vertices, and $Q^M$ is the vector of the blood flow rate of net outflow from vertices. By splitting inner vertex and boundary vertex-related terms into different sides of the equation, the linear system can be transformed to[12]:

$$G_{int}^M \cdot P_{int}^M = G_b^M \cdot P_b^M$$

$G_{int}^M$ is the submatrix of the hydraulic conductance matrix that contains inner vertex rows, while $G_b^M$ only contains boundary vertex rows. With given boundary blood pressure values $P_b^M$, the unknown inner vertex blood pressure $P_{int}^M$ can be effectively solved using the generalized minimum residual method (GMRES)[13]

#### Phase Separation Effect

Red blood cells (RBC) have a highly heterogeneous distribution in real vasculature, especially in poorly structured tumor vasculatures[14], due to the nonproportional distribution of RBCs in daughter branches at diverging bifurcations known as the phase separation effect[15]. It is important to obtain an estimation of RBC distribution in vasculature not only because of its effect on apparent blood viscosity but also because of its dominating role in oxygen delivery. Approximately 98% of the oxygen carried in the blood is bound to hemoglobin contained in RBCs, while only 2% is dissolved in plasma and RBC water[16]. Vessels without adequate red blood cells are unable to support the tissue metabolism despite the flow rate.

In this study, we assume a uniform discharge hematocrit feed of 0.45[17] at the vasculature inlet. In the downstream bifurcation vertices in the vasculature, we adopted an experimentally determined parametric description of the phase separation effect by Pries AR et al.[7]:

$$logit(FQ_E) = A + B \cdot logit\left(\frac{FQ_B - X_0}{1 - 2X_0}\right)$$

$$A = -13.29 \cdot \frac{{D_\alpha}^2 / {D_\beta}^2 - 1}{{D_\alpha}^2 / {D_\beta}^2 + 1} \cdot \frac{1 - H_d}{D_p}$$

$$B = 1 + 6.98 \cdot \frac{1 - H_d}{D_p}$$

$$X_0 = 0.964 \cdot \frac{1 - H_d}{D_p}$$

$D_p$, $D_\alpha$ and $D_\beta$ are the vessel diameters of the parent vessel and two daughter vessels measured in microns. $H_d$ is the discharge hematocrit (Hd) of the parent vessel. $FQ_E$ is defined as the fractional flow of RBCs into the daughter branch $\alpha$ and $FQ_B$ is the corresponding blood flow fraction. The RBC distribution at each bifurcation-vertex can be calculated as shown above using a bisection method. Combining this with a depth-first search (DFS)-like algorithm, the $Hd$ distribution in the entire vasculature network can be obtained efficiently.

#### Iterative Update

Discharge hematocrit levels significantly influence the apparent viscosity of blood in each vessel segment, potentially altering the flow patterns within the vasculature system. To tackle the interaction between blood flow and discharge hematocrit distribution, we employ an iterative approach. This involves solving a linear system for blood flow and conducting vasculature traversal for discharge hematocrit until the system reaches convergence. Typically, with a physiologically plausible vasculature structure, convergence is achieved with low iteration differences within 3-4 iterations.

#### Edge Contraction

The node-tube representation of vasculature data effectively captures vascular morphology and vessel shape. However, the extensive number of nodes and tubes could significantly complicate the computation of blood flow. To improve computational efficiency, we developed a corresponding vertex-edge model specifically for hemodynamic calculations within the vasculature. This approach leverages the fact that non-leaking edges exhibit consistent flow through all constituent tubes, meaning that only the aggregate information of each edge and the pressure at its vertices are required to assess perfusion. By applying edge contraction to the vasculature, conducting hemodynamic computations on this equivalent vertex-edge model, and then mapping the results back to the original node-tube representation, we significantly enhance computational efficiency, achieving improvements by orders of magnitude.

#### S2.2 Initialization and boundary conditions

##### Oxygen Dynamics in Host Tissue

Despite the high degree of microvascular heterogeneity arising from both extrinsic and intrinsic factors, control mechanisms in healthy tissue effectively mitigate variations in oxygen supply, resulting in a relatively stable and adequate oxygen concentration[18]. This observation is supported by studies such as that by Carreau et al.[19], which report low spatial variation in oxygen concentration in various healthy organs, including the brain, muscle, and intestinal tissue. In light of these findings, and to avoid the need to explicitly model the healthy tissue vasculature, we proposed a dynamic equilibrium oxygen partial pressure distribution in healthy tissue governed by:

$$\frac{\partial P_{oxy}}{\partial t} = \nabla \cdot \left(D_{oxy} \nabla P_{oxy}\right) - \frac{1}{a_{oxy}} M\left(P_{oxy}\right) + \frac{1}{a_{oxy}} S_h\left(P_{boxy} - P_{oxy}\right)$$

$S_h$ represents the oxygen supply from healthy tissue vasculature, the value is selected such that the steady state oxygen concentration in normal tissue equals $P_{hs}$ :

$$M(P_{hs}) = S_h\left(P_{boxy} - P_{hs}\right)$$

##### Host Vasculature Initialization

The vascularization through angiogenesis requires pre-existing host vasculature to start with. Common methods establishing host vasculature includes the cubic grid vasculature[20], parallel vessels arrays[21], and reduced vasculature data containing a few major vessels[22]. However, existing methods can hardly fulfill our need for unbiased tumor growth. The artificial vessel arranged as cubic grid or paralleled lines could introduce bias in certain growth direction while the density of reduced vasculature could be inadequate for nutrition supply and angiogenesis initiation. Ideally, the entire vasculature extracted from normal tissue is preferred for host vasculature initialization, however, with the scarcity of data availability and the cost of handing a much larger simulation region and vasculature size, makes it currently impractical. As an alternative approach, we propose a novel spatially stratified tangent vessel method for host vasculature initialization. This method provides a simple and theoretically unbiased vascular environment for tumor development and allows for random sampling to further annihilation the risk of directional growth bias introduced error in statistical analysis.

In our model, the initial tumor assumes a spherical shape with a radius $R_{Tumor}^{init} = 150\ \mu\text{m}$. The environmental vasculature surrounding the tumor is organized as tangent lines on an extended sphere situated $L_{S2V}^{init} = 50\ \mu\text{m}$ away from the tumor's surface. Both the tangent points and vessel orientations on the tangent plane are randomly sampled. To mitigate the risk of large vessel-starved areas, which are not physiologically reasonable in a healthy host, we implement a stratification strategy. This involves sampling the tangent points part by part within uniformly divided subregions on the surface of the sphere. Each initialized vessel is 6 μm in radius, with a total length of $L_{HostVes}^{init} = 1000\ \mu\text{m}$.

#### Boundary Blood Pressure

The boundary blood pressure assignment is also crucial for proper perfusion and the downstream vessel remodeling estimation. For a static system, convex optimization method considering the WSS under a mass conservation constraint[6] could be used to estimate the boundary blood pressure. However, the optimization-based estimation is impractically expensive for an evolving vasculature. Tailored to our dynamic development task and the specific host vasculature morphology, we introduce a novel location-encoded blood pressure assignment method to determine the inlet and outlet blood pressures of the vasculature under significant deformation. This method incorporates two key components.

**Distance-to-center-based baseline pressure term**: This term adjusts the blood pressure based on the distance from the vessel center. This ensures a progressive increase in pressure difference between the inlet and outlet as the vessel length extends.

**Angular-position-based pressure variation term**: This term introduces a blood pressure shift for host vessels with different orientation and facilitates the establishment of appropriate pressure gradients within neo-vasculatures connecting different host vessels, especially in the central region.

For the location-encoded blood pressure assignment method, the initial step involves estimating the vessel length based on the position of the inlet or outlet. We assume that throughout the near-spheroidal tumor growth, for the host vessel ends, their distance to tumor surface and angular position with respect to tumor center remains constant. And for other parts in host vasculature, their distance to tumor surface does not drop below initial surface to vessel distance $L_{S2V}^{init}$. Based on these assumptions, we introduce another geometric approximation to decompose any host vessel into three components: one circular arc component on the extended sphere surface, and two tangent line components connecting the vessel ends and the arc.

The constant host vessel to tumor surface distance and the constant angle between two ends of a host vessel with respect to the tumor center throughout the tumor development is:

$$L_{S2E} = \sqrt{\left(R_{Tumor}^{init} + L_{S2V}^{init}\right)^2 + \left(\frac{L_{HostVes}^{init}}{2}\right)^2} - R_{Tumor}^{init}$$

$$\theta_{ECE} = 2arctan\left(\frac{L_{HostVes}^{init}}{2\left(R_{Tumor}^{init} + L_{S2V}^{init}\right)}\right)$$

Then as the tumor grows, the estimated distance from tumor center to the arc sphere at time t is:

$$L_{C2V}^{t} = L_{C2E}^{t} - L_{S2E} + L_{S2V}^{init}$$

Where $L_{C2E}^{t}$ represents the distance from the evaluated vessel end to the tumor center at time t, which is the only required variable. The angle between vessel end and the corresponding tangent point is:

$$\theta_{ECT}^{t} = \arccos\left(\frac{L_{C2V}^{t}}{L_{C2E}^{t}}\right)$$

And the total length of the deformed host vessel is written as:

$$L_{vas}^{t} = 2L_{C2E}^{t}(\theta_{ECE} - \theta_{ECT}^{t} + \sin(\theta_{ECT}^{t}))$$

Finally, the baseline pressure term is:

$$P_{D2C}^{t}(L_{vas}^{t}) = \pm\, \kappa_{D2C}\frac{L_{vas}^{t}}{2} + P_{ref}$$

Where $P_{ref}$ is the reference mean blood pressure of the vasculature, $\kappa_{D2C}$ is the gradient for distance-based blood pressure term, which is determined according to the reference WSS and radius of the vasculature. Sign in the first term determines the flow direction which is randomly assigned during initialization with a positive gradient for inlet and a negative for outlet.

The angular position of the vessel end can be calculated as:

$$\theta_{C2E}^{t} = arccos(\widehat{\boldsymbol{n}}_{C2E}^{t} \cdot \widehat{\boldsymbol{z}})$$

$$\varphi_{C2E}^{t} = arctan\left(\frac{\widehat{\boldsymbol{n}}_{C2E}^{t} \cdot \widehat{\boldsymbol{y}}}{\widehat{\boldsymbol{n}}_{C2E}^{t} \cdot \widehat{\boldsymbol{x}}}\right)$$

Where $\widehat{\boldsymbol{n}}_{C2E}^{t}$ is the unit vector pointing from tumor center to the vessel end, θ is the polar angle and ϕ is the azimuthal angle. $\widehat{\boldsymbol{x}}$, $\widehat{\boldsymbol{y}}$, and $\widehat{\boldsymbol{z}}$ are unit axis vectors of the cartesian coordinate system.

The angular-position-based pressure term is:

$$P_{\text{angle}}^{t} = c_{angle}\kappa_{D2C}\frac{L_{C2E}^{t}}{\sqrt{2}}[\cos(2\theta_{C2E}^{t}) + \sin(2\theta_{C2E}^{t})\sin(2\varphi_{C2E}^{t})]$$

Where $c_{angle}$ represents the relative strength of angular variation compared to the baseline gradient term. This low-frequency angular term ensures that vessel ends positioned at approximately opposite angles share similar pressure variation values. This guarantees approximately constant blood pressure gradients for all host vessels throughout their growth.

#### S2.3 Model Parameters

##### Continuum model parameters

| Parameter | Description | Value | Unit | Reference |
|---|---|---|---|---|
| $\boldsymbol{\phi_{Host}}$ | Blatz-Ko model parameter | 1 | | Richard Moran et al.[23] |
| $\boldsymbol{\beta_{Host}}$ | Blatz-Ko model parameter | 2 | | Richard Moran et al.[23] |
| $\boldsymbol{\mu_{Host}}$ | Host shear modulus | 1 | $kPa$ | Richard Moran et al.[23] |
| $\boldsymbol{\phi_{Tumor}}$ | Blatz-Ko model parameter | 0.2 | | James MacLaurin[24] |
| $\boldsymbol{\beta_{Tumor}}$ | Blatz-Ko model parameter | 4 | | James MacLaurin[24] |
| $\boldsymbol{\mu_{Tumor}}$ | Tumor shear modulus | 2.7 | $kPa$ | James MacLaurin[24] |
| $\boldsymbol{P_N}$ | Critical $P_{oxy}$ for necrosis | 0.1 | $mmHg$ | |
| $\boldsymbol{\rho}$ | Tissue density | 1 | $^{g}/_{ml}$ | |

| | | | | |
|---|---|---|---|---|
| $\boldsymbol{K_g}$ | Growth rate | 0.693 | $day^{-1}$ | See text |
| $\boldsymbol{P_{\lambda 50}}$ | $P_{oxy}$ at half maximum growth rate | 10 | $mmHg$ | See text |

*Tabel S3 Biophysical parameters used in the tumor growth model.*

#### Discrete model parameters

| Module | Parameter | Description | Value | Unit | Reference |
|---|---|---|---|---|---|
| Oxygen | $D_{oxy}$ | Oxygen diffusion coefficient | 2410 | $\mu m^2 s^{-1}$ | Bentley et al.[25] |
| | $a_{oxy}$ | Oxygen solubility | 38.9 | $nlO_2 ml^{-1} mmHg^{-1}$ | Bentley et al.[25] |
| | $\rho$ | Tissue mass density | 1000 | $kg \cdot m^{-3}$ | |
| | $M_{max}$ | Max oxygen consumption | 2-4 | $mmHg \cdot s^{-1}$ | See text |
| | $P_{M50}$ | $P_{oxy}$ with half - maximum consumption | 1 | $mmHg$ | Goldman[26] |
| | $P_{boxy}$ | Blood $P_{oxy}$ | 35.5 | $mmHg$ | |
| | $P_{hs}$ | Steady state host tissue $P_{oxy}$ | 35 | $mmHg$ | Carreau et al.[19] |
| | $P_{cap}$ | Max oxygen pressure difference across vessel wall | 1 | $mmHg$ | |
| Perfusion | $\mu_0$ | Apparent viscosity of rat blood at 37℃ | 1.05 | $mPa \cdot s$ | Pries et al.[7] |
| | $P_{ref}$ | Reference mean blood pressure | 40 | $mmHg$ | |
| | $\kappa_{D2C}$ | Gradient for distance-based blood pressure term | 1.8752 | $mmHg \cdot mm^{-1}$ | |
| | $c_{angle}$ | Relative strength of angular blood pressure variation | 2 | | |
| TAF | $C_{max}$ | Max TAF concentration | 1 | | |
| | $D_{TAF}$ | TAF Diffusion coefficient | 20 | $\mu m^2 s^{-1}$ | Alberding et al.[27] |

| | | | | | |
|---|---|---|---|---|---|
| | $K_{TAF}$ | TAF decay rate | 0.002 | $s^{-1}$ | Adapted from Alberding et al.[27] |
| | $P_{TAF}$ | Tissue $P_{oxy}$ where cells start to release TAF | 34.5 | $mmHg$ | |
| **Angiogenesis** | $k_{sprout}$ | Maximum sprout rate per length | 0.05 | $\mu m^{-1} day^{-1}$ | |
| | $C_{TAF50}$ | $C_{TAF}$ for half-maximal sprout rate | 0.33 | | |
| | $C_{th}$ | $C_{TAF}$ threshold for sprout formation | 0.01 | | |
| | $C_{mig}$ | $C_{TAF}$ threshold for stalk cell proliferation | 0.01 | | |
| | $R_{sprout}$ | Radius of sprout | 6 | $\mu m$ | |
| | $V_{sprout}$ | Velocity of sprout elongation | 75 | $\mu m \cdot day^{-1}$ | Levi B. Wood et al.[28] |
| | $k_{TAF}$ | Weight for TAF gradient | 1 | | |
| | $k_{ana}$ | Weight for anastomosis bias | 1 | | |
| | $k_{rand}$ | Weight for random variation | 0.5 | | |
| | $D_{ana}$ | Maximum tip cell sensing distance | 75 | $\mu m$ | Gerhardt [29] |
| | $\theta_{ana}$ | Maximum vessel sensing angle | $\pi/3$ | | Secomb et al.[30] |
| | $L_{ana}$ | Anastomosis threshold | 25 | $\mu m$ | |
| **Remodeling** | $v_{bamax}$ | Branching angle remodeling velocity | 1 | $\mu m \cdot h^{-1}$ | |
| | $F_{bath}$ | Branching angle remodeling threshold | 0.25 | | JP Alberding et al.[27] |
| | $T_s$ | Structural adaptation coefficient | 192 | $day$ | |
| | $n_r$ | Adaptation weighting factor for reference radius | 4 | | |

| | | | | | |
|---|---|---|---|---|---|
| | $R_{ref}$ | Reference radius in adaptation | 12 | $\mu m$ | |
| | $\tau_{ref}$ | Reference wall shear stress | 15 | $dyn \cdot cm^{-2}$ | Secomb et al.[6] |
| **Initialization** | $R_{Tumor}^{init}$ | Initial tumor radius | 150 | $\mu m$ | |
| | $L_{S2V}^{init}$ | Initial tumor surface to host vasculature distance | 50 | $\mu m$ | |
| | $L_{HostVes}^{init}$ | Initial host vasculature length | 1000 | $\mu m$ | |
| | $N_{HostVes}^{init}$ | Initial host vessel number | 50 | | |
| | $L_{Host}^{init}$ | Initial host tissue cube length | 4000 | $\mu m$ | |

*Table S4 Biophysical Parameters related to vasculature and oxygenation.*
